# Revisiting the role of mean annual precipitation in shaping functional trait distributions at a continental scale

**DOI:** 10.1101/2023.06.29.546983

**Authors:** Isaac R. Towers, Peter A. Vesk, Elizabeth H. Wenk, Rachael V. Gallagher, Saras M. Windecker, Ian J. Wright, Daniel S. Falster

**Affiliations:** Evolution & Ecology Research Centre, University of New South Wales Sydney, Australia; School of Agriculture, Food and Ecosystem Sciences, The University of Melbourne, Parkville, Australia; Hawkesbury Institute for the Environment, Western Sydney University, Penrith, Australia

**Keywords:** mean annual precipitation, continental-scale, functional trait, climate, dynamic global vegetation model, plant growth form, woodiness

## Abstract

Mean annual precipitation (MAP) plays an undisputed role in determining the spatial distribution of the vegetative ecosystems on Earth. Nevertheless, the relationship between MAP and plant functional traits remains unclear. Here, we test the relationship between eight key functional traits and MAP. Our analysis reveals a strong, coordinated response of several plant traits including leaf mass per area, leaf nitrogen, the leaf carbon isotope ratio and plant height from resource-conservative to resource-acquisitive values as MAP increased. These results establish an important role for MAP in driving trait selection across space and, therefore, a need for these effects to be included in future theoretical frameworks.

## 2 Introduction

Mean annual precipitation (MAP) varies substantially across the globe, impacting the spatial distribution and structure of vegetation (Schimper, 1898). However, evidence for consistent relationships between MAP and the functional traits of the organisms in these ecosystems is equivocal. Some early empirical analyses reported MAP as a key predictor of plant traits (Moles *et al*., 2009; Wright *et al*., 2005; Ordoñez *et al*., 2009), but more recent global analyses found weaker relationships. For example, Maire *et al*. (2015) showed soil variables were more influential predictors than climate for most analysed traits, Moles *et al*. (2014) found MAP to be a weaker predictor of trait values than mean annual temperature (MAT), and Bruelheide *et al*. (2018) found that MAP, among other broad-scale environmental variables, was a poor predictor of community-weighted mean trait values. The role of MAP in shaping the spatial distribution of species traits, and thus the effect of MAP on key ecosystem processes, has therefore remained unclear.

It has recently been proposed that regionalising the scale of analysis might reveal trait-environment patterns in greater clarity, by controlling for factors such as phylogenetic history (Kambach *et al*., 2023). Here, we revisit our understanding of the relationship between traits and MAP using an unprecedented database, AusTraits - the largest harmonised continent-specific collection of georeferenced trait values globally. Australia represents the ideal laboratory to test trait-MAP relationships for several reasons. Firstly, MAP and MAT are orthogonal in Australia (r = 0.04; Figure 1a). As such, independent associations between water availability and traits can be isolated from the effect of MAT. Australia also spans an extraordinary precipitation gradient, encompassing the 22nd (82 mm/year) and 99th (4211 mm/year) quantiles of the global distribution of MAP, thereby representing all but the very driest regions of the globe (Figure 1b). Consequently, trait data can be readily obtained from a range of biomes including arid and semi-arid regions which are typically not well represented in global analyses. Finally, although Australia is a major land-based carbon sink (accounting for approximately 60% of the global terrestrial carbon sink in some years (Poulter *et al*., 2014)), there is also significant uncertainty regarding the effect of precipitation on carbon uptake in this region (Teckentrup *et al*., 2021). This uncertainty is proposed to emerge from a number of factors including 1) disagreement amongst dynamic vegetation models (DVMs) regarding the processes governing plant response to water limitation, 2) poor representation of drought-adaptation within the highly endemic and structurally distinct vegetation that is present in Australia, and 3) significant variation within model ensembles in the simulated or prescribed fraction of woody and herbaceous cover (Teckentrup *et al*., 2021). Altogether, a re-examination of the relationship between plant traits and MAP would not only improve our fundamental understanding of the evolution of trait distributions but would yield a timely assessment of the embedded processes in DVGMs used to simulate ecosystem processes (Teckentrup *et al*., 2021).

**Figure 1:**
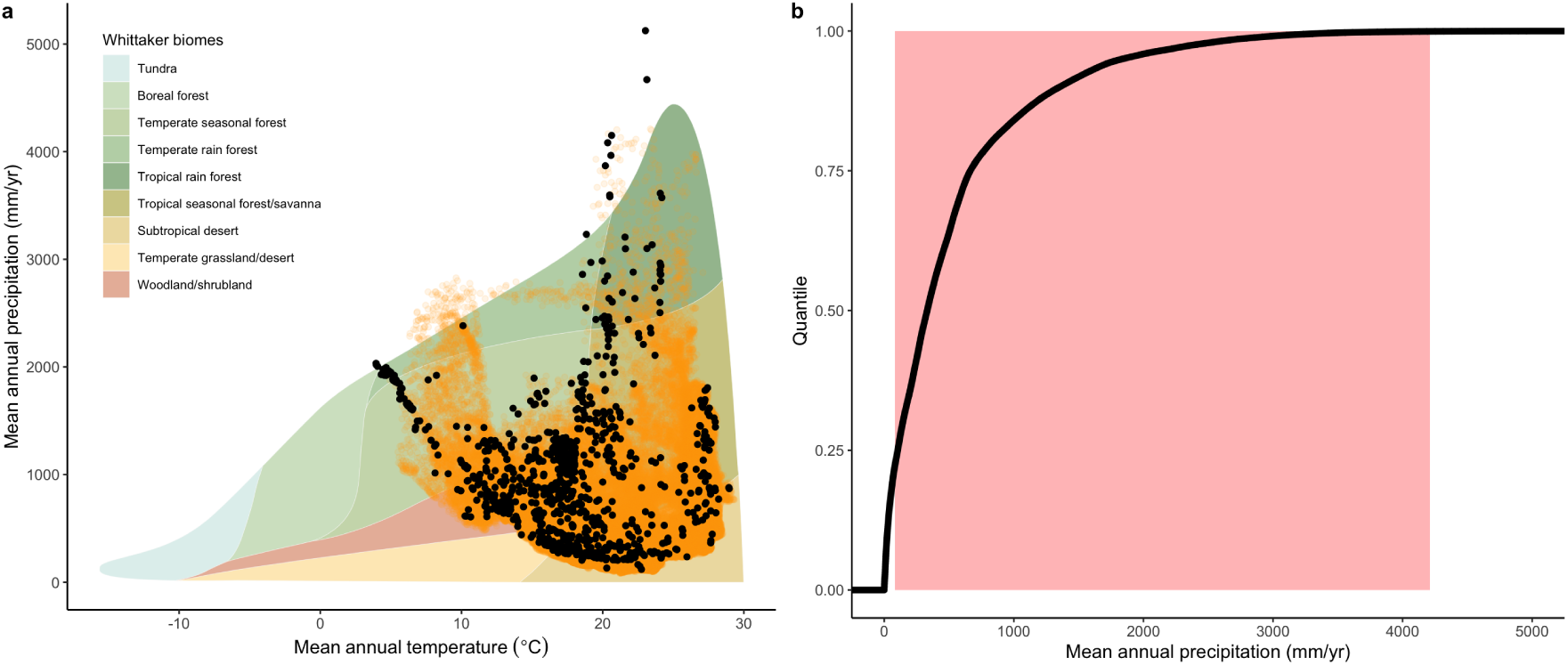
Comparison of the climate space of Australia relative to the global climate space. In panel **a**, the location of sites where the focal traits for the present study were sampled (black points) and the Australian climate space (orange points) are overlaid on the Whittaker biome classification system which classifies biomes according to mean annual precipitation (mm) and mean annual temperature (*°C*). Panel **b** is the empirical cumulative density functions of the global climate space for mean annual precipitation. The red polygon indicates the span of the Australian climate.

We selected eight key functional traits widely considered to capture important physiological processes in vascular plants and generated hypotheses for how each would respond to spatial variation in MAP. These were derived from published eco-evolutionary theories (Table 1) explicitly relating traits to MAP or soil moisture and, if these were not available, we inferred predictions from theories based on other moisture-related environmental drivers including VPD and site productivity. We used bivariate linear regressions to test each of these hypotheses and inferences (according to the sign of the relationship). To account for potential variation in trait responses to the environment due to woodiness, we tested whether observed patterns observed differed when species were classified as woody or non-woody. We expected relationships would be stronger in woody taxa because long-lived individuals must function in challenging environmental conditions, whereas non-woody species often avoid dry conditions by surviving as seed.

**Table 1:**
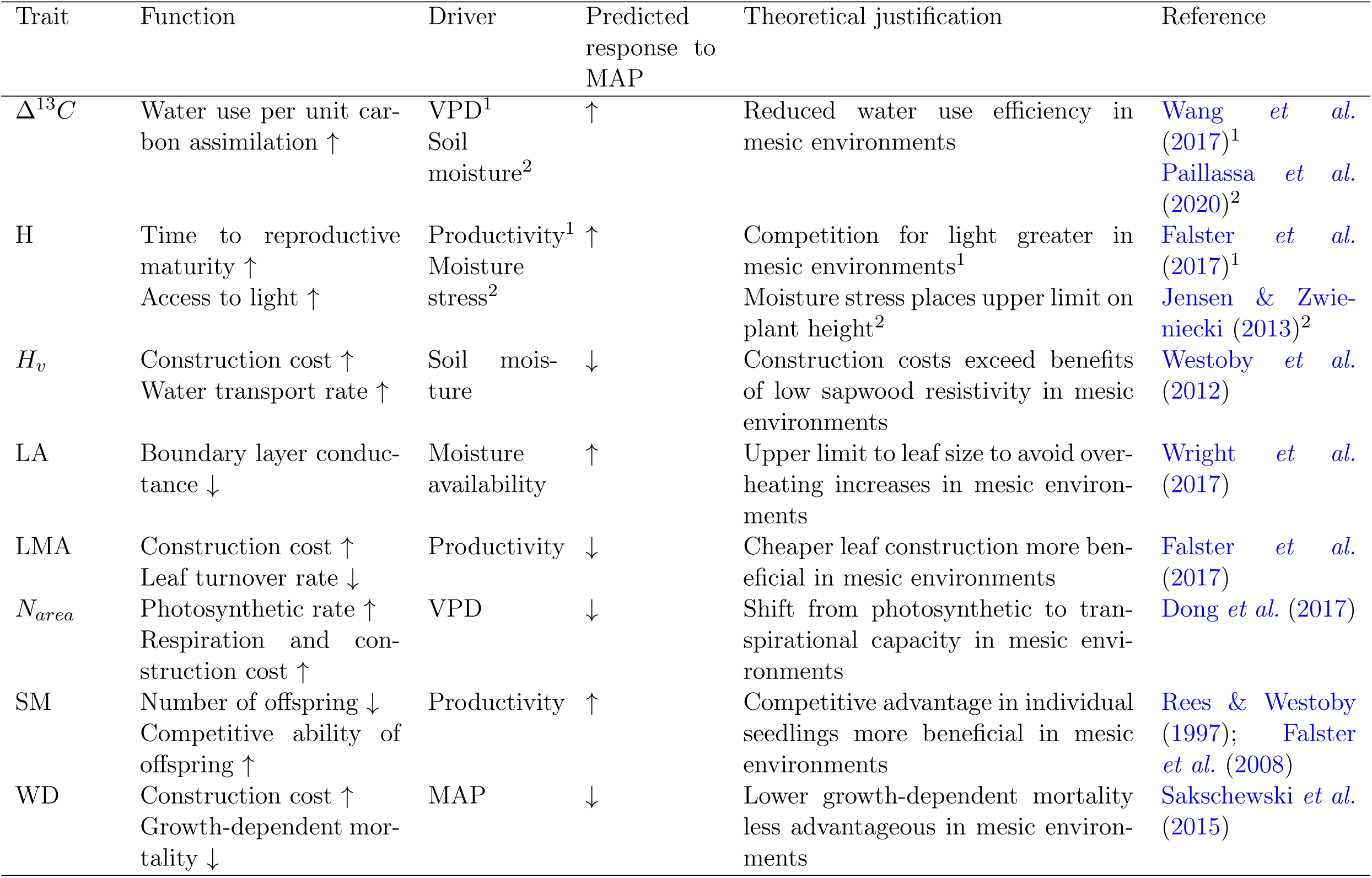
The functional description of each trait, its predicted response to MAP based on its theorised response to a given environmental driver in the literature and the theoretical justification for the prediction. Arrows in "Function" indicate the effect of an increase in the trait value on the listed function while arrows in "Predicted response to MAP" indicate the predicted direction of the trait response to increasing MAP. Superscripts indicate that theoretical expectations for plant height are derived from different sources

## 3 Results

For woody taxa, MAP was an excellent predictor (i.e. *r*^2^ *≥* 30%) of the variation in Δ^13^*C* (C3 plants only), leaf mass per area (LMA), plant height (H) and leaf nitrogen per-area (*N_area_*), with Δ^13^*C* and LMA decreasing and H and *N_area_*increasing with MAP (Table 2; Figure 2). In addition, MAP was a moderate predictor of leaf area (LA; *r*^2^ = 0.23), being positively correlated with MAP. However, MAP was a relatively weak predictor (i.e. *r*^2^ *≤* 20%) of wood density (WD) and Huber value (*H_v_*), being slightly negatively correlated, and was a poor predictor of seed mass (SM; *r*^2^ *∼* 12%). Regardless of correlation strength, in all cases the direction of the fitted correlation was consistent with our predictions (Table 1).

**Figure 2:**
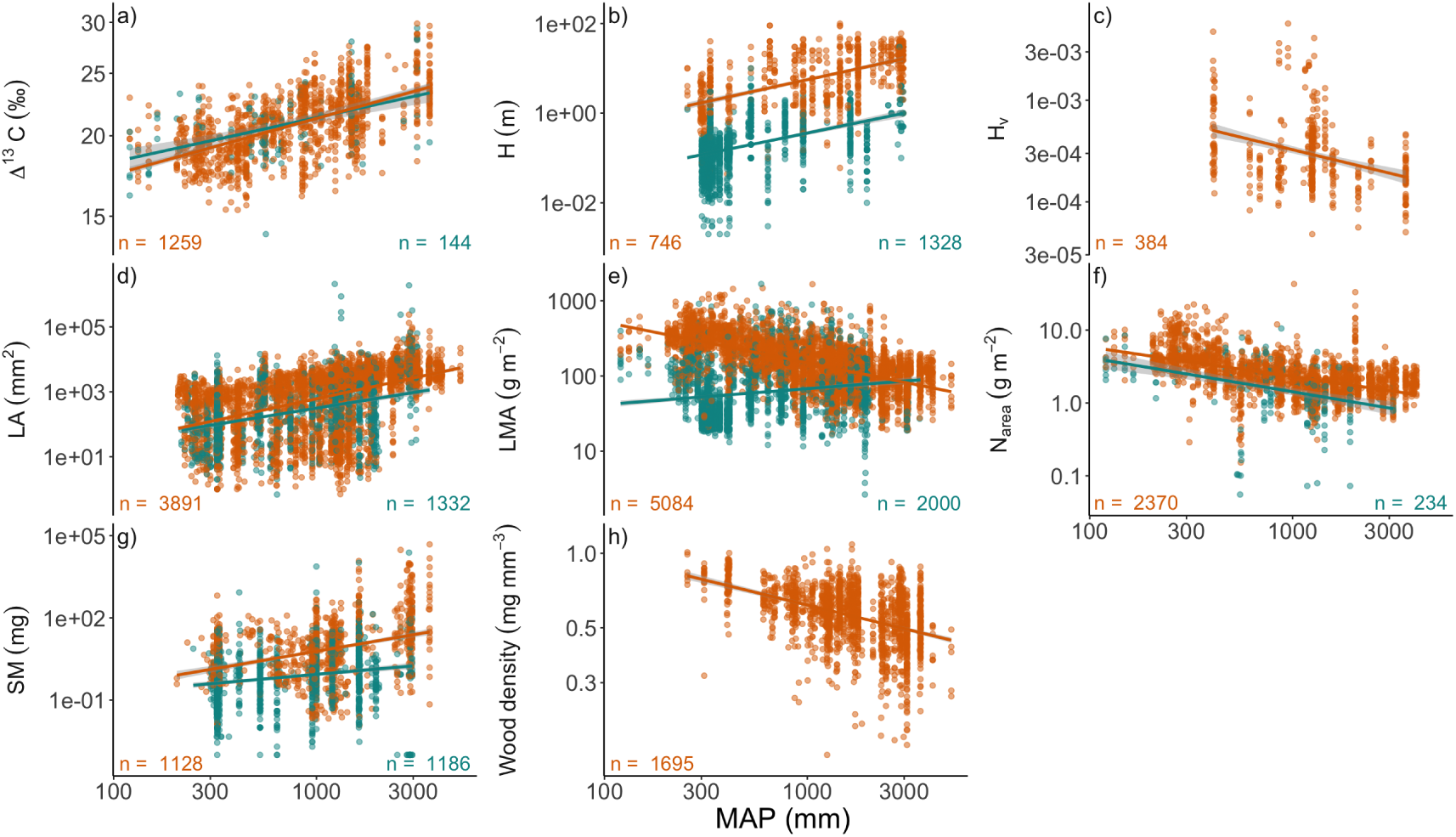
Relationships between eight functional traits and mean annual precipitation (MAP). Observations are species-by-site mean values. Orange circles are woody taxa and teal circles are non-woody taxa. Solid lines are the fitted curves from linear regression while the grey fields represent the 95% confidence interval around the predicted curve. Note the log-scaled axes. *r*^2^ for each relationship are available in Table 2.

**Table 2:** Variance in each functional trait (*r*^2^) explained by the linear relationship with mean annual precipitation when all taxa are analyzed together (overall) and when separated by growth form. Sign indicates the direction of the relationship. Analyses of the wood-related traits wood density (WD) and huber value (*H_v_*) were not conducted for non-woody taxa. The direction of all relationships were consistent with theoretical expectations for woody taxa. Number of observations is available in Figure 2. All traits are log-transformed.

| Trait | Overall |  | Woody |  | Non-woody |  |
| --- | --- | --- | --- | --- | --- | --- |
| | $r^2$ | Sign | $r^2$ | Sign | $r^2$ | Sign |
| $\Delta^{13}C$ | 0.35 | + | 0.36 | + | 0.30 | + |
| H | 0.42 | + | 0.32 | + | 0.18 | + |
| $H_v$ | | | 0.14 | - | | |
| LA | 0.24 | + | 0.23 | + | 0.15 | + |
| LMA | 0.02 | - | 0.38 | - | 0.03 | + |
| $N_{area}$ | 0.25 | - | 0.30 | - | 0.17 | - |
| SM | 0.13 | + | 0.12 | + | 0.04 | + |
| WD |  |  | 0.19 | - |  |  |

Combining woody and non-woody observations had opposing effects on the amount of variance explained by MAP for the six non-wood related traits (see Methods). Specifically, we found a substantial increase in the variation explained for H (*r*^2^ = 42%) accompanied by a minor increase in the variation explained for SM (*r*^2^ = 0.13%). However, there was a notable reduction in the variance explained for *N_area_*, and a dramatic decrease in the variation of LMA explained, to the extent that MAP and LMA were now largely uncorrelated. These outcomes emerged because, for the most part, trait-MAP relationships were weaker in non-woody taxa and, in the case of LMA, there was a bimodal distribution in the observations at low MAP due to a greater representation of herbaceous annuals.

## 4 Discussion

In contrast to recent global analyses (Maire *et al*., 2015; Moles *et al*., 2014; Bruelheide *et al*., 2018), we found MAP was a strong predictor of several key functional traits, in a manner consistent with predictions based on the theoretical literature (Table 1). Specifically, as MAP increased we observed a coordinated shift from resource-conservative to resource-acquisitive values of most traits. This response was most prominent in woody taxa, in partial support of our hypothesis, but we also found important exceptions where the explanatory power of MAP was equivalent between growth forms. These findings have important consequences for the theoretical underpinnings of trait evolution in response to broad-scale environmental gradients and, in an applied context, the development of encoded relationships which describe how vegetation interacts with precipitation in the next generation of trait-based DVMs.

The strongest trait-MAP relationship we observed was the positive correlation between plant height and MAP when all taxa were analysed simultaneously. This outcome is qualitatively similar to a previous global-scale study of plant height, which found that precipitation in the wettest quarter, followed closely by MAP, were the strongest predictors of plant height (Moles *et al*., 2009). In contrast to that study, however, we found that MAP explained a substantially greater amount of the total variation in the data (40% compared to 21%). Given the considerable spread in plant heights that exists within sites (Figure 2), the fact that MAP explained over 40% of the variation suggests that it is a major driver of the site-to-site variation in plant stature.

Plant height may increase with MAP for several reasons. Maximum achievable height may be biophysically constrained by water availability such that, all else being equal, taller plants can survive in wetter sites (Jensen & Zwieniecki, 2013). While this is a plausible explanation, we also observed that particularly tall species occurred across almost the entire MAP gradient (Figure 2). As such, an alternate explanation is that natural selection favours taller plants, on average, in wetter sites because the benefit of having a higher position in the canopy outweighs the drawback of increased stem construction costs and delayed reproduction as competition for light intensifies (Falster *et al*., 2017).

Similar to plant height, we also found that leaf Δ^13^*C* and MAP were strongly and positively correlated across all taxa. Δ^13^*C* is a measure of the long-term average of the ratio of the partial pressure of CO_2_ in the intercellular spaces (*C_i_*) and the atmosphere (*C_a_*). Consistent with a recent, global-scale empirical analysis (Cornwell *et al*., 2018), we found a strong positive effect of MAP on Δ^13^*C*. According to least-cost theory, all else being equal, the cost of procuring water for transpiration increases in drier environments (Prentice *et al*., 2014), leading to reduced stomatal conductance, lower *C_i_*, and causing plants to invest more in photosynthetic capacity relative to water transport capacity (Dong *et al*., 2017). Indeed, in line with this theory, we also observed that *N_area_*, which is linked to photosynthetic capacity (Dong *et al*., 2022), was greater in sites with lower MAP regardless of growth form. Remarkably, in contrast to Cornwell *et al*. (2018), the strength of the correlation between Δ^13^*C* and MAP was strong for both woody and non-woody taxa, suggesting that the processes governing stomatal behaviour are similarly sensitive to MAP in both growth forms. The analysis of Δ^13^*C* presented here, however, was limited to C3 species owing to the well understood link between Δ^13^*C* and the *C_i_* : *C_a_* ratio in these taxa.

Separately analysing the trait data for woody and non-woody species also revealed important differences in response to MAP. For example, in line with our expectations based on theory, LMA was found to be strongly and negatively correlated with MAP (i.e. *r*^2^ *∼* 40%) for woody taxa. LMA is theorised to decline as site productivity increases because the more rapid height growth rates conferred by cheaply-constructed leaves, and therefore greater access to light, compensates for high leaf turnover costs associated with the lower leaf life-span of low LMA leaves (Falster *et al*., 2017). In addition, higher LMA is hypothesised to be advantageous in more arid sites to maintain leaf function at more negative leaf water potentials (Wang *et al*., 2023). Qualitatively, our results are consistent with Wright *et al*. (2004), who found a modest negative relationship (i.e. *r*^2^ *∼* 10%) between LMA and MAP at the global-scale, but only after statistically controlling for MAT. The fact that MAP explained such a large proportion of the variation in LMA in the current study relative to the global-scale is intriguing. In Australia, woody trees and shrubs are mostly evergreen, meaning that the well-recognised trade-off between LMA and leaf life span is not obscured by a shift to being cold-deciduous. Even after analysing the data separately for evergreen versus deciduous taxa, however, Wright *et al*. (2004) found MAP only explained at most 22% of the variation in LMA. The observed relationship points to an important role of MAP in driving selection on leaf economic strategies for woody taxa across strong rainfall gradients in Australia, and perhaps beyond.

In contrast, LMA and MAP were virtually uncorrelated when analysing just the non-woody taxa and this reflected the general tendency for trait-environment relationships to be weaker within this functional group. Trait interdependencies, or trait trade-offs, which yield the relationships we observed in woody taxa, may not be present in the non-woody taxa analysed. For example, an analysis of trait connectivity using data from the TRY database revealed that the direct relationship between leaf life-span and LMA which is seen across all taxa and woody taxa is not present for non-woody taxa (Flores-Moreno *et al*., 2019). Alternatively, weaker trait-environment relationships may emerge in non-woody species because this functional group employs a wider variety of strategies to tolerate harsher or more variable environments, such as occupying relatively benign positions in the understorey or employing dormancy as a bet-hedging strategy in the seed bank as is the case for many annual herbs analysed (Dwyer & Erickson, 2016).

### 4.1 Vegetation modelling and future directions

Increasing reliability in DVMs has been accompanied by the development of trait-based approaches which allow trait values to be predicted directly based on presiding climatic conditions (De Kauwe *et al*., 2015). Our analysis using a harmonised, data-rich trait database provides clear, albeit correlation-based, evidence for a role of MAP in shaping broad-scale trait gradients in Australia and, consequently, impetus for these relationships to be incorporated into the next generation of DVMs. However, this integration depends on our ability to mechanistically describe the processes that cause these relationships to emerge. To this end, we note that the traits with the strongest correlations with MAP (i.e. LMA, H, *N_area_*) tended to lack theoretical hypotheses directly linking them to precipitation or soil moisture, being driven instead by site productivity or VPD (Table 1).

The simple analysis that we conducted here also invites further investigation of trait-environment correlations using this unprecedented dataset for Australian taxa. For instance, our treatment of species-by-site combinations as independent data points, although suitable for analysing mean trends, precludes investigation of the relative contribution of species turnover versus within-species variation emerging from phenotypic plasticity or local adaptation to the observed patterns (Ackerly & Cornwell, 2007). Identifying the sources of this variation would have important implications for our understanding of how trait-climate patterns have emerged historically, and how resilient communities will be to future change (Dong *et al*., 2020). In a similar manner, the substantial variation that we observed for some traits across species within sites points to a significant contribution of local-scale species coexistence to trait diversity, the magnitude of which may itself be dependent on climate (Andrew *et al*., 2021). For example, although we observed a mean negative response of WD to MAP, there was also a strong triangular signal in the data where taxa in mesic sites had a much greater range of WD from very light to very dense wood (Figure 2). Substantial coexistence of strategies within a given climatic band could also go some way to explain the very weak response of SM to precipitation that we and others (Moles *et al*., 2005) have observed. The challenge for theoretical frameworks, therefore, is not just to predict the central tendency of trait distributions (Dong *et al*., 2017; Wang *et al*., 2023; Xu *et al*., 2021; Wang *et al*., 2017), but also the likely diversity of strategies which may occur under a given climatic regime. Theoretical models which incorporate biotic interactions among multiple, potentially co-existing strategies into the process of selection (i.e. game-theoretic models) seem most promising in this regard.

## 5 Conclusion

Using simple linear correlations, and guided by quantitative predictions and inferences from the literature, we have demonstrated that for long-lived, woody taxa, MAP is strongly coordinated with key functional traits. Although the scope of this study was limited to Australia, we propose that the underlying mechanism driving the observed patterns are widely generalisable, but are potentially obscured by distinct phylogenetic history, endemism and/or covariation with other environmental factors Gallagher *et al*. (2023). To this end, we advocate for re-focusing the geographical scope of analyses to regions where observations span strong gradients in the variable of interest independently of other environmental covariates (Kambach *et al*., 2023).

## 6 Acknowledgments

Discussions with Will Cornwell, Mark Westoby, David Coleman and John Dwyer enriched the analysis and interpretation of the results. This research was funded by a grant awarded by Eucalypt Australia (Eucalypt futures: using functional traits to predict species distributions and responses to environmental change) to DSF and PAV.

## 8 Author contributions

IRT and DSF conceived the original idea with further development of the concept by PAV, EHW, RVG, SMW and IJW. IRT conducted the review of trait theory. IRT conducted statistical analysis with assistance from SMW. IRT led the writing of the manuscript. DSF, PAV, EHW, RVG, SMW and IJW assisted with drafting the manuscript. All authors approved of the final draft of the manuscript.

## 9 Data availability

The data and code used to support the findings in the manuscript will be made openly available on Zenodo if accepted.

## 11 Methods

### 11.1 Trait data

AusTraits (Version 4.1.0; Falster *et al*. (2021)) is a harmonised trait database containing over 1,000,000 individual records of functional traits for over 30,000 taxa occurring in Australia (including offshore islands and territories). It is therefore the largest collation of trait data for plants growing in Australia and for any whole continent.

We queried the AusTraits database for eight functional traits which represent multiple dimensions of plant ecophysiological strategy, are widely distributed across Australia (Figure S1,S2,S3,S4), and for which predictions about the effect of MAP could be derived from the literature (Table 1). These traits were the leaf carbon isotope ratio (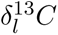), plant height (H), huber value (*H_v_*), leaf area (LA), leaf mass per area (LMA), leaf nitrogen per area (*N_area_*), dry seed mass (SM) and wood density (WD). Because we were interested in how traits respond to variation in climate, we filtered the data to include only observations which had associated geospatial information. In order to sample "natural" trait observations from AusTraits, we also limited the scope of our study to trait observations made on unmanipulated individuals occurring *in situ*. In other words, this excluded most data originating from field, laboratory and glasshouse experiments, herbarium records, and expert opinion.

After applying the above criteria, we also queried the AusTraits database for studies which did not have *N_area_* values but instead reported values for leaf nitrogen per dry mass (*N_mass_*) and LMA for the same individuals, which could be converted into *N_area_* using the following equation:

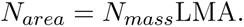

This process yielded an additional 3170 individual-level observations of *N_area_*. We also converted 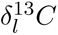 (‰) into the inferred leaf carbon isotope discrimination value (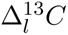; ‰) using the following equation:

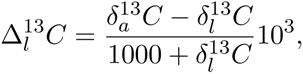

where the free atmospheric carbon isotope ratio, 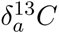 (‰), is -8 (Farquhar *et al*., 1989).

Measurement protocols for LA on compound species differed between studies which considered LA at the leaflet versus the whole-leaf scale. The theoretical framework for predicting the response of LA to the environment considered in the present study is based on the rate of conductance through the leaf-atmosphere boundary layer which, for compound species, is likely determined at the leaflet-scale (Wright *et al*., 2017). Thus, to minimise variation in the LA data associated with the varying scale of measurement, we first identified datasets which measured LA on compound taxa and for which LA was measured at the leaf scale, or for which the measurement protocol was unknown. Then, within these datasets only, we used the "leaf_compoundness" data in AusTraits, which classifies taxa as being "simple", "compound", or "simple compound", to group taxa as having either simple or compound leaves, assuming "simple compound" taxa to be compound. LA observations for taxa with compound leaves, or for which compoundness was unknown within these datasets were then omitted. In sum, approximately 2000 of the 17,500 independent LA records were removed after this process.

We used a number of approaches to check for errors in the data. Firstly, to assess for dataset-level errors, we compared the data distribution for a given dataset against the distribution of all other datasets combined for each trait and visually inspected this comparison for obvious discrepancies. Then, for taxa with multiple observations, especially those with observations from different contributors, we inspected various metrics for each trait including the ratio of the maximum and minimum value of the trait, the coefficient of variation and the absolute values of the minimum and maximum to identify potentially erroneous observations. Although we attempted to correct data points when possible (usually owing to unit entry), we removed a number of data points, typically being those which had unusually high or low values relative to other observations of that taxa but also to the distribution of the trait overall. Finally, we inspected the data for duplication of trait values for a given taxon between contributors, which may occur in cases where the data was obtained from a shared data source.

To address our hypothesis regarding the influence of plant growth form on the relationship between MAP and functional traits, we also queried the AusTraits database for the "woodiness_detailed" trait, using a dataset compiled by Wenk (*in progress*). This trait describes taxa based on both the presence and vertical extent of secondary xylem (i.e. "true wood") and thus seperates taxa into multiple categories. In the case of this study, we did not view plant growth form as a response variable *per se*. Instead, we pooled taxa with growth form information into two possible groups being either "woody" or "non-woody". For the "woody" group, we selected taxa with a erect habit and a lignified stem based on the "woody" category in the "woodiness_detailed" dataset. This category was the single largest group in the "woodiness_detailed" dataset, capturing just over half of the taxa for which information was available. By comparison, the non-woody group captured a more diverse group of growth forms, including herbs (being the most dominant non-woody group) but also tussocks, graminoids and palmoid taxa among other classifications. Some taxonomic groups such as monocots and ferns can occasionally produce lignified wood-like material despite lacking secondary xylem. To test whether classification of these taxa into a given group influenced the outcome of our analysis, we conducted our analysis twice, including taxa with the definition "woody_like_stem" first into the "non-woody" and secondly into the "woody" group. There was a minor increase in model performance for both the woody and non-woody group in the latter case, so we primarily report the results of this set of analyses, although there was no qualitative difference in the interpretation of our results in either case (Table S1,S2,S3,S4).

### 11.2 Climate data

Although our specific aim was to investigate the relationship between traits and MAP, we also selected a range of other climatic variables which are considered to describe soil and atmospheric moisture availability. Climate data were obtained from a variety of sources, as no single source provided all variables. Mean annual precipitation, mean precipitation in the wettest quarter, mean precipitation in the driest quarter and mean precipitation seasonality were obtained from Worldclim at a resolution of 1/120° (*∼* 1*km*), mean annual potential evapotranspiration and mean annual Thornwaite aridity index from Envirem at a resolution of 1/120° and mean monthly VPD data from 1981-2010 from CHELSA at a resolution of 1/120° (*∼* 4*km*). For the mean monthly VPD, we averaged the values across months to find a mean annual VPD. We then extracted climate data corresponding to the geospatial information associated with trait observations. In peninsular coastal regions where trait observations were recorded, climate data were occasionally not available. In these cases, we used the *nearestLand* function from the *seegSDM* package Golding & Shearer (2023) to assign these observations with climate data from the nearest non-NA cell within 10 km. If this correction still did not yield climate data, these observations were not included in statistical analyses.

Overall, observed relationships tended to be quantitatively similar or weaker when replacing MAP with other variables (Table S5, S2, S4). The most notable exceptions were that there was some evidence to suggest that *N_area_* responded more negatively to MI and LA more positively to precipitation in the wettest quarter and seasonality compared to MAP in both woody and non-woody taxa when separated as well as when combined. Thus, although we report these results in the Supplementery Information, we did not focus on these findings.

### 11.3 Statistical analysis

Our analyses considered that trait variation may occur at two spatial scales being within-site variation and across-site variation. Here, a site is the scale across which broad-scale climatic gradients influence trait variation. Variation within sites is instead assumed to be due to other factors including within-site environmental heterogeneity as well as local coexistence of multiple functional strategies. Here, we assigned trait observations to sites by overlaying a 1/120° grid on the continent, corresponding to the resolution of the climate data, where each grid cells represents a site. Prior to statistical analysis, we then found the site-level mean for each taxon for each trait, yielding 400-5000 unique species-by-site combinations depending on the trait. In other words, for a given trait, each site has a number of observations equivalent to the number of taxa recorded in it. This means that species-by-site combinations sometimes averaged across observations from different data contributors and through time.

Prior to statistical analysis, we transformed trait and climate data to meet the assumption of normality for linear regression. Specifically, we log-transformed all traits, and log-transformed VPD, MAP and precipitation seasonality.

To first test trait responses to MAP across all available taxa, we regressed the mean species-by-site values of each functional trait against MAP using ordinary least-squares regression. We then assessed the Pearson correlation coefficient as a measure of the strength and direction of correlations as well as the *r*^2^ value to understand the degree of model fit. Given the large sample sizes involved in this study, p-values for even weak correlations are likely to be significant so we do not report them. To assess the explanatory power of MAP compared to other moisture-related variables, we regressed each trait as a bivariate linear function of each of the remaining climatic variables. To test our hypothesis that woody taxa will exhibit stronger responses to the climate than non-woody taxa, we then repeated the process as described above for the woody and non-woody groups separately. In all cases, taxa with unknown growth forms were omitted from analyses to permit comparison between patterns when taxa were combined and when separated by growth form.

To explore the possibility of non-linear curvature in the relationship between traits and climate, we fit all of the above models again with an additional quadratic parameter, and assessed the *r*^2^ value to identify whether including the quadratic term significantly improved model fit. For the most part, model fits were not improved substantially (Table S6,S7,S8,S9,S10). There was, however, some evidence that *N_area_* had a concave-up relationship with MAP whereby there was a steep decline in *N_area_* from arid to mesic regions until *∼*1200mm rainfall where it began increasing again slightly (Figure 2).

## 13 Supplementary information

**Table S1:**
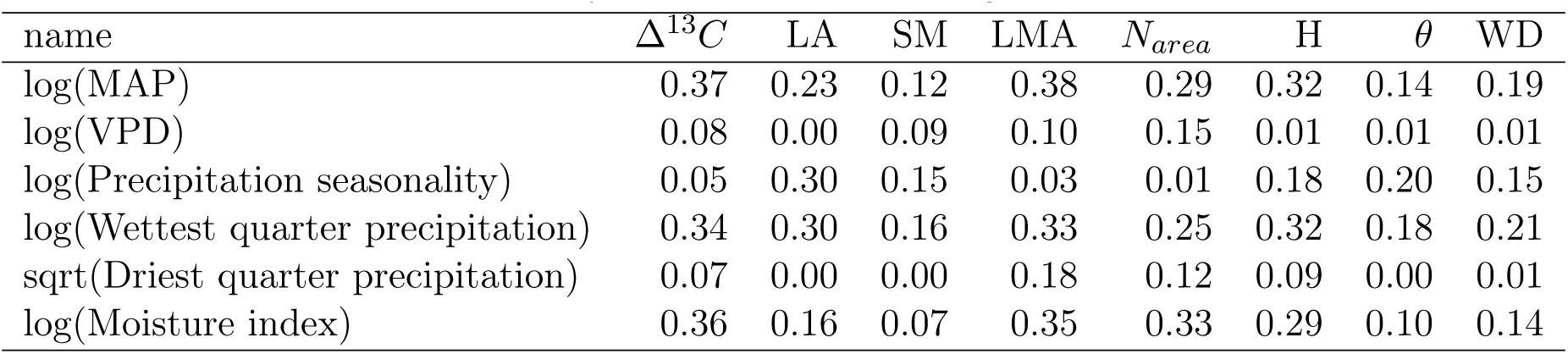
*r*^2^ values for the linear relationship between each functional trait and environmental variable for woody taxa. All traits are log-transformed.

| name | $\Delta^{13}C$ | LA | SM | LMA | $N_{area}$ | H | $\theta$ | WD |
| --- | --- | --- | --- | --- | --- | --- | --- | --- |
| log(MAP) | 0.37 | 0.23 | 0.12 | 0.38 | 0.29 | 0.32 | 0.14 | 0.19 |
| log(VPD) | 0.08 | 0.00 | 0.09 | 0.10 | 0.15 | 0.01 | 0.01 | 0.01 |
| log(Precipitation seasonality) | 0.05 | 0.30 | 0.15 | 0.03 | 0.01 | 0.18 | 0.20 | 0.15 |
| log(Wettest quarter precipitation) | 0.34 | 0.30 | 0.16 | 0.33 | 0.25 | 0.32 | 0.18 | 0.21 |
| sqrt(Driest quarter precipitation) | 0.07 | 0.00 | 0.00 | 0.18 | 0.12 | 0.09 | 0.00 | 0.01 |
| log(Moisture index) | 0.36 | 0.16 | 0.07 | 0.35 | 0.33 | 0.29 | 0.10 | 0.14 |

**Table S2:** *r*^2^ values for the linear relationship between each functional trait and environmental variable for woody taxa and woody-like taxa without secondary xylem. All traits are log-transformed.

| name | $\Delta^{13}C$ | LA | SM | LMA | $N_{area}$ | H | $\theta$ | WD |
| --- | --- | --- | --- | --- | --- | --- | --- | --- |
| log(MAP) | 0.36 | 0.23 | 0.12 | 0.38 | 0.30 | 0.32 | 0.14 | 0.19 |
| log(VPD) | 0.08 | 0.00 | 0.09 | 0.10 | 0.15 | 0.01 | 0.01 | 0.01 |
| log(Precipitation seasonality) | 0.05 | 0.30 | 0.15 | 0.03 | 0.01 | 0.18 | 0.20 | 0.15 |
| log(Wettest quarter precipitation) | 0.33 | 0.30 | 0.16 | 0.33 | 0.25 | 0.32 | 0.18 | 0.21 |
| sqrt(Driest quarter precipitation) | 0.07 | 0.00 | 0.00 | 0.18 | 0.12 | 0.09 | 0.00 | 0.01 |
| log(Moisture index) | 0.36 | 0.16 | 0.07 | 0.35 | 0.33 | 0.29 | 0.10 | 0.14 |

**Table S3:** *r*^2^ values for the linear relationship between each functional trait and environmental variable for non-woody taxa. All traits are log-transformed.

| name | $\Delta^{13}C$ | LA | SM | LMA | $N_{area}$ | H |
| --- | --- | --- | --- | --- | --- | --- |
| log(MAP) | 0.27 | 0.15 | 0.04 | 0.03 | 0.16 | 0.18 |
| log(VPD) | 0.01 | 0.00 | 0.01 | 0.01 | 0.20 | 0.04 |
| log(Precipitation seasonality) | 0.16 | 0.16 | 0.03 | 0.09 | 0.00 | 0.13 |
| log(Wettest quarter precipitation) | 0.30 | 0.19 | 0.05 | 0.01 | 0.14 | 0.12 |
| sqrt(Driest quarter precipitation) | 0.00 | 0.03 | 0.01 | 0.03 | 0.14 | 0.17 |
| log(Moisture index) | 0.22 | 0.10 | 0.02 | 0.03 | 0.20 | 0.15 |

**Table S4:** *r*^2^ values for the linear relationship between each functional trait and environmental variable for non-woody taxa when woody-like taxa without secondary xylem are not included in analysis. All traits are log-transformed.

| name | $\Delta^{13}C$ | LA | SM | LMA | $N_{area}$ | H |
| --- | --- | --- | --- | --- | --- | --- |
| log(MAP) | 0.30 | 0.15 | 0.04 | 0.03 | 0.17 | 0.18 |
| log(VPD) | 0.00 | 0.00 | 0.01 | 0.01 | 0.21 | 0.04 |
| log(Precipitation seasonality) | 0.17 | 0.16 | 0.03 | 0.10 | 0.00 | 0.13 |
| log(Wettest quarter precipitation) | 0.32 | 0.19 | 0.05 | 0.01 | 0.14 | 0.12 |
| sqrt(Driest quarter precipitation) | 0.00 | 0.03 | 0.01 | 0.04 | 0.14 | 0.17 |
| log(Moisture index) | 0.25 | 0.10 | 0.02 | 0.03 | 0.21 | 0.15 |

**Table S5:**
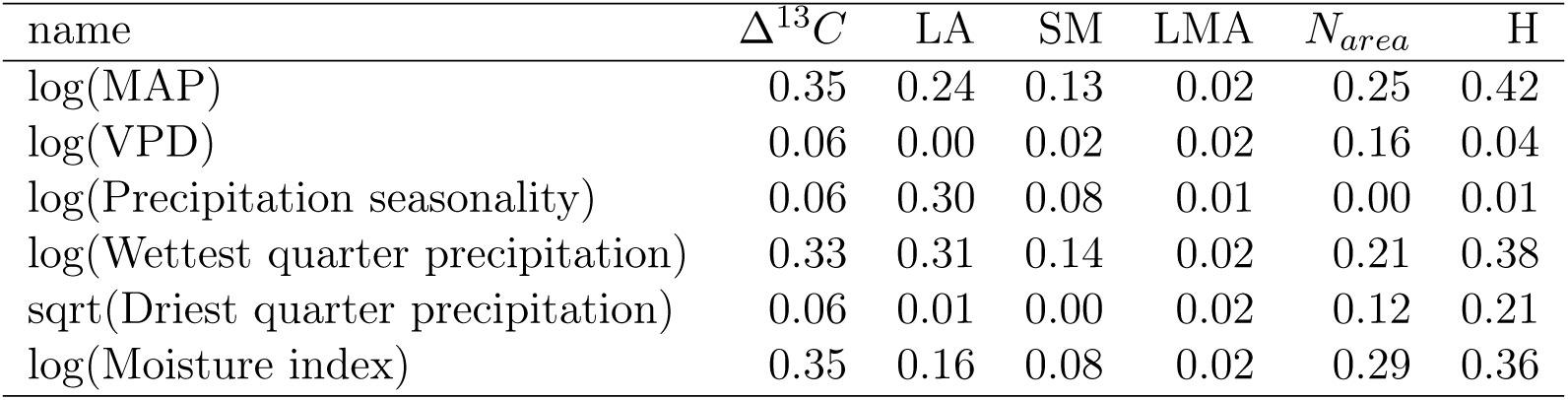
*r*^2^ values for the linear relationship between each functional trait and environmental variable for all taxa. All traits are log-transformed.

| name | $\Delta^{13}C$ | LA | SM | LMA | $N_{area}$ | H |
| --- | --- | --- | --- | --- | --- | --- |
| log(MAP) | 0.35 | 0.24 | 0.13 | 0.02 | 0.25 | 0.42 |
| log(VPD) | 0.06 | 0.00 | 0.02 | 0.02 | 0.16 | 0.04 |
| log(Precipitation seasonality) | 0.06 | 0.30 | 0.08 | 0.01 | 0.00 | 0.01 |
| log(Wettest quarter precipitation) | 0.33 | 0.31 | 0.14 | 0.02 | 0.21 | 0.38 |
| sqrt(Driest quarter precipitation) | 0.06 | 0.01 | 0.00 | 0.02 | 0.12 | 0.21 |
| log(Moisture index) | 0.35 | 0.16 | 0.08 | 0.02 | 0.29 | 0.36 |

**Table S6:** *r*^2^ values for the quadratic relationship between each functional trait and environmental variable for all taxa. All traits are log-transformed.

| name | $\Delta^{13}C$ | LA | SM | LMA | $N_{area}$ | H |
| --- | --- | --- | --- | --- | --- | --- |
| log(MAP) | 0.36 | 0.26 | 0.14 | 0.03 | 0.29 | 0.42 |
| log(VPD) | 0.07 | 0.06 | 0.03 | 0.05 | 0.17 | 0.05 |
| log(Precipitation seasonality) | 0.06 | 0.30 | 0.09 | 0.03 | 0.01 | 0.16 |
| log(Wettest quarter precipitation) | 0.33 | 0.33 | 0.15 | 0.02 | 0.25 | 0.40 |
| sqrt(Driest quarter precipitation) | 0.25 | 0.01 | 0.02 | 0.02 | 0.16 | 0.27 |
| log(Moisture index) | 0.36 | 0.16 | 0.08 | 0.02 | 0.31 | 0.36 |

**Table S7:** *r*^2^ values for the quadratic relationship between each functional trait and environmental variable for woody taxa. All traits are log-transformed.

| name | $\Delta^{13}C$ | LA | SM | LMA | $N_{area}$ | H | $\theta$ | WD |
| --- | --- | --- | --- | --- | --- | --- | --- | --- |
| log(MAP) | 0.37 | 0.26 | 0.18 | 0.38 | 0.35 | 0.32 | 0.14 | 0.19 |
| log(VPD) | 0.08 | 0.04 | 0.10 | 0.15 | 0.15 | 0.03 | 0.01 | 0.03 |
| log(Precipitation seasonality) | 0.05 | 0.30 | 0.15 | 0.04 | 0.01 | 0.18 | 0.20 | 0.16 |
| log(Wettest quarter precipitation) | 0.34 | 0.32 | 0.21 | 0.33 | 0.29 | 0.32 | 0.19 | 0.21 |
| sqrt(Driest quarter precipitation) | 0.25 | 0.01 | 0.05 | 0.18 | 0.16 | 0.09 | 0.01 | 0.02 |
| log(Moisture index) | 0.38 | 0.16 | 0.12 | 0.35 | 0.37 | 0.29 | 0.10 | 0.14 |

**Table S8:** *r*^2^ values for the quadratic relationship between each functional trait and environmental variable for woody taxa and woody-like taxa without secondary xylem. All traits are log-transformed.

| name | $\Delta^{13}C$ | LA | SM | LMA | $N_{area}$ | H | $\theta$ | WD |
| --- | --- | --- | --- | --- | --- | --- | --- | --- |
| log(MAP) | 0.37 | 0.26 | 0.18 | 0.38 | 0.35 | 0.32 | 0.14 | 0.19 |
| log(VPD) | 0.08 | 0.04 | 0.10 | 0.15 | 0.15 | 0.03 | 0.01 | 0.03 |
| log(Precipitation seasonality) | 0.05 | 0.30 | 0.16 | 0.04 | 0.01 | 0.18 | 0.20 | 0.16 |
| log(Wettest quarter precipitation) | 0.33 | 0.32 | 0.21 | 0.33 | 0.29 | 0.32 | 0.19 | 0.21 |
| sqrt(Driest quarter precipitation) | 0.25 | 0.01 | 0.06 | 0.18 | 0.16 | 0.09 | 0.01 | 0.02 |
| log(Moisture index) | 0.38 | 0.16 | 0.12 | 0.35 | 0.37 | 0.29 | 0.10 | 0.14 |

**Table S9:** *r*^2^ values for the quadratic relationship between each functional trait and environmental variable for non-woody taxa. All traits are log-transformed.

| name | $\Delta^{13}C$ | LA | SM | LMA | $N_{area}$ | H |
| --- | --- | --- | --- | --- | --- | --- |
| log(MAP) | 0.29 | 0.16 | 0.05 | 0.03 | 0.17 | 0.20 |
| log(VPD) | 0.02 | 0.07 | 0.01 | 0.01 | 0.20 | 0.04 |
| log(Precipitation seasonality) | 0.17 | 0.17 | 0.06 | 0.10 | 0.01 | 0.16 |
| log(Wettest quarter precipitation) | 0.31 | 0.21 | 0.05 | 0.02 | 0.16 | 0.12 |
| sqrt(Driest quarter precipitation) | 0.27 | 0.05 | 0.03 | 0.05 | 0.16 | 0.23 |
| log(Moisture index) | 0.23 | 0.10 | 0.04 | 0.04 | 0.21 | 0.17 |

**Table S10:** *r*^2^ values for the quadratic relationship between each functional trait and environmental variable for non-woody taxa when woody-like taxa without secondary xylem are not included in analysis. All traits are log-transformed.

| name | $\Delta^{13}C$ | LA | SM | LMA | $N_{area}$ | H |
| --- | --- | --- | --- | --- | --- | --- |
| log(MAP) | 0.32 | 0.16 | 0.04 | 0.03 | 0.17 | 0.20 |
| log(VPD) | 0.02 | 0.07 | 0.01 | 0.01 | 0.21 | 0.04 |
| log(Precipitation seasonality) | 0.18 | 0.17 | 0.06 | 0.11 | 0.02 | 0.16 |
| log(Wettest quarter precipitation) | 0.34 | 0.21 | 0.05 | 0.02 | 0.17 | 0.12 |
| sqrt(Driest quarter precipitation) | 0.29 | 0.05 | 0.03 | 0.05 | 0.15 | 0.23 |
| log(Moisture index) | 0.26 | 0.10 | 0.04 | 0.03 | 0.21 | 0.17 |

**Figure S1:**
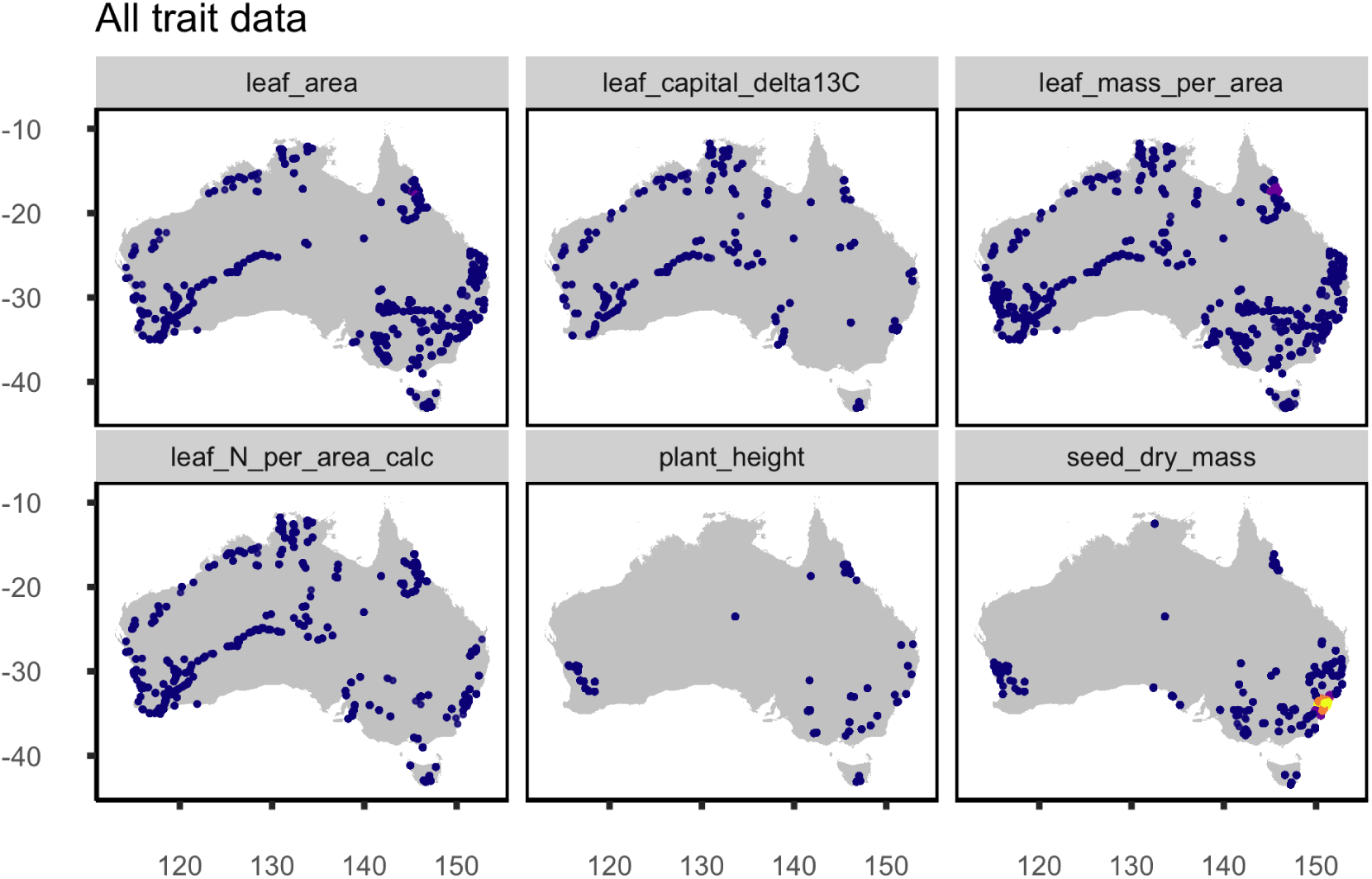
Geographical distribution of data for eight functional traits for all taxa in Aus-Traits. Blue to yellow colour gradient for points represents increasing number of overlapping points.

**Figure S2:**
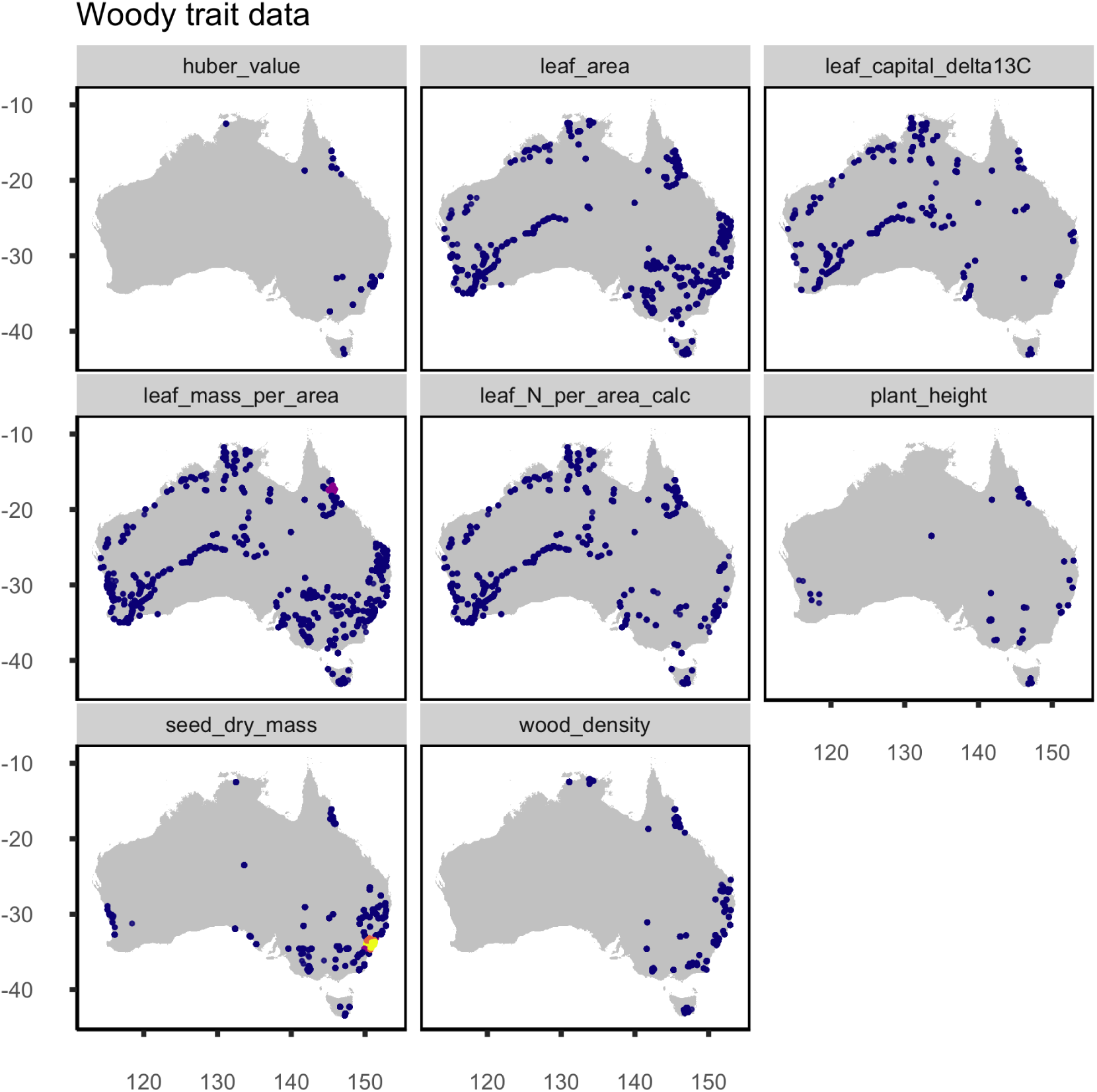
Geographical distribution of data for eight functional traits for woody taxa only in AusTraits. Point colour as in Figure S1.

**Figure S3:**
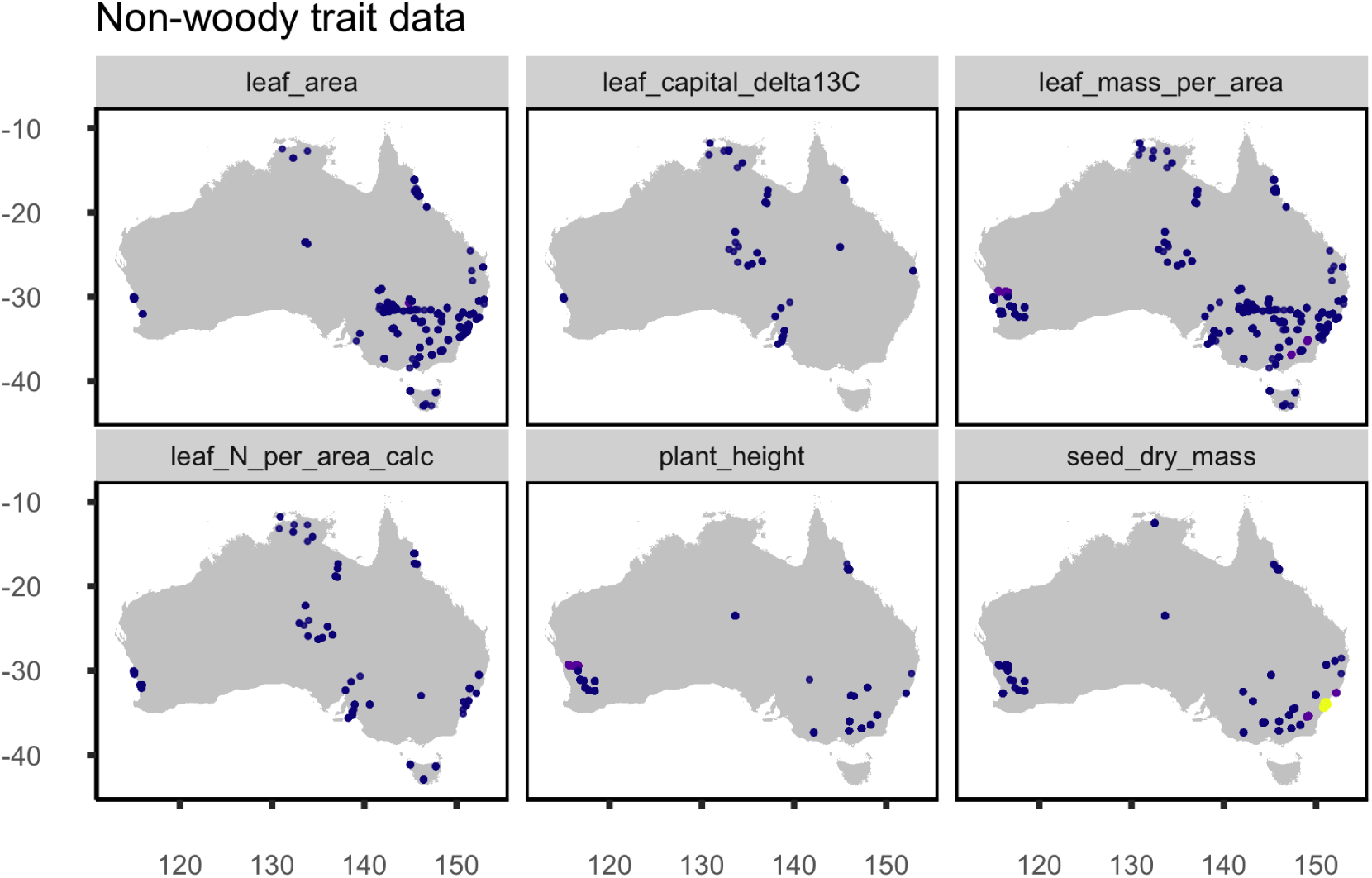
Geographical distribution of data for eight functional traits for non-woody taxa only in AusTraits. Point colour as in Figure S1.

**Figure S4:**
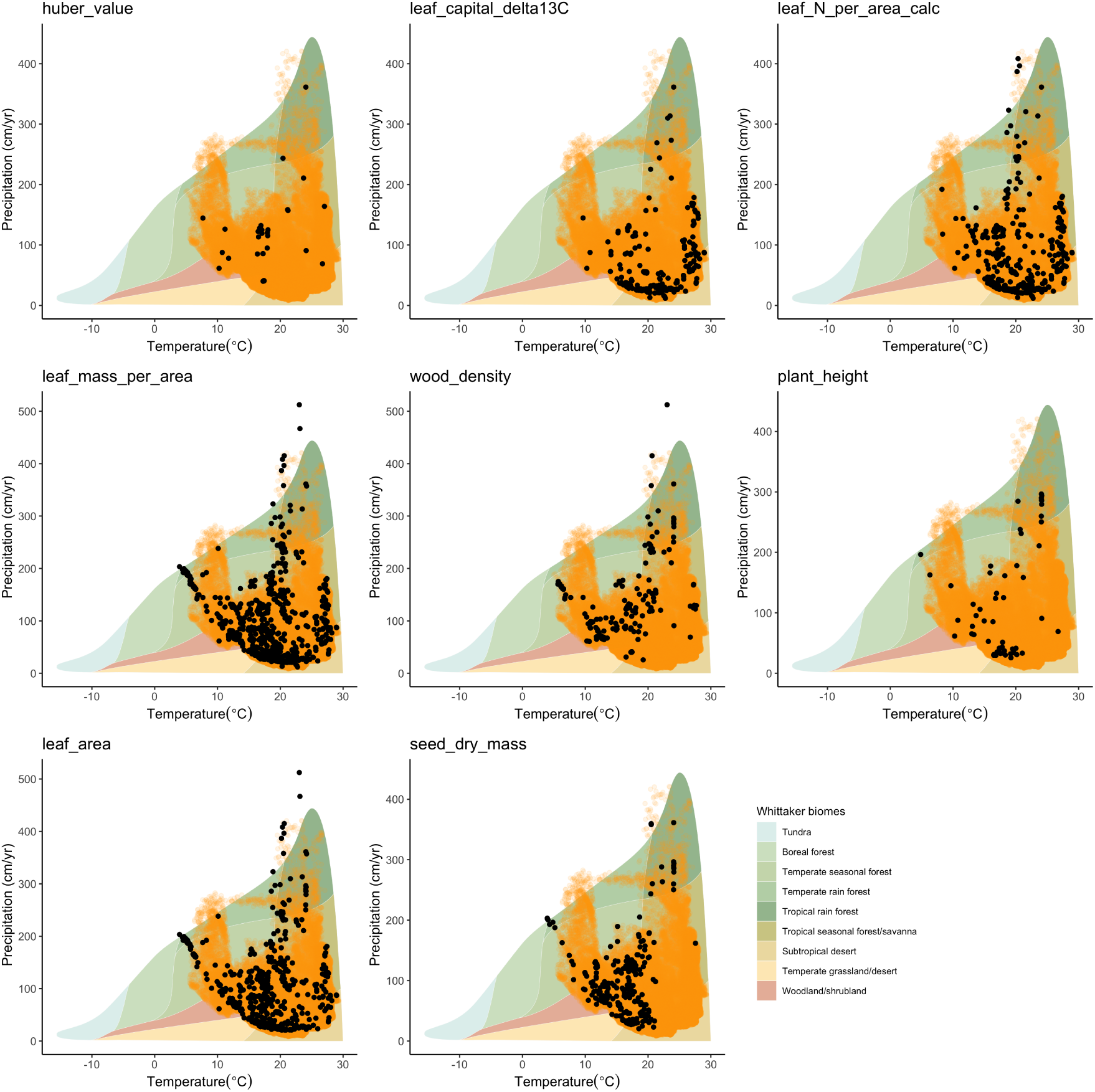
Distribution of trait observations in AusTraits across Whittaker’s biomes with respect to mean annual precipitation (cm/yr) and mean annual temperature (°C). Orange points are the climatic values present on the Australian continent and the black points are trait observations.

